# Effect of acute ultraviolet radiation on *Galleria mellonella* health and immunity

**DOI:** 10.1101/2022.12.02.517514

**Authors:** Ausrine Sabockyte, Samuel McAllister, Christopher J. Coates, Jenson Lim

## Abstract

For humans, acute and chronic overexposure to ultraviolet (UV) radiation can cause tissue damage in the form of sunburn and promote cancer(s). The immune-modulating properties of UV radiation and health-related consequences are not well known. Herein, we used the larvae of the wax moth (*Galleria mellonella*) to determine UV-driven changes in cellular components of innate immunity. From immune cell (haemocyte) reactivity and the production of antimicrobial factors, these insects share many functional similarities with mammalian innate immunity. After exposing insects to UVA or UVB, we monitored larval viability, susceptibility to infection, haemolymph (blood) physiology and faecal discharge. Prolonged exposure of larvae to UVB coincided with decreased survival, enhanced susceptibility to bacterial challenge, melanin synthesis in the haemolymph, compromised haemocyte functionality and changes in faecal (bacterial) content. We contend *G. mellonella* is a reliable *in vivo* model for assessing the impact of UV exposure at the whole organism and cellular levels.

## INTRODUCTION

Ultraviolet (UV) light is one of the components of sunlight and consists of UVA (400-320nm), UVB (320-290nm) and UVC (290-200nm) and has both beneficial and detrimental effects on human health either through the skin, eye or immune system (D’Orazio et al, 2013). Although UV light represents an important source of non-ionising radiation that induces the synthesis of Vitamin D, which modulates the immune system (Hart et al, 2011), and can also act against the immune system by inhibiting antigen presentation, stimulating the release of immunosuppressive cytokines and induces the generation of lymphocytes of the regulatory subtype (Schwarz, 2002). The major molecular target for UV-induced immunosuppression is DNA damage as the radiation is absorbed by adjacent pyrimidine DNA bases and cis-urocanic acid, which leads to the production of reactive oxygen species (ROS) causing DNA damage, mutations in oncogenes, and the onset of cellular carcinogenesis (Halliday 2005; González Maglio, 2016). In mammals, this can lead to immunosuppression by regulating the levels of pro- and anti-inflammatory cytokines and/or the induction of regulatory T cells (Sreevidya et al, 2010; Garibyan et al, 2010; Schwarz et al, 2010; Schwarz et al, 2011. However, cells possess mechanisms of protection against DNA damage, an example being melanocytes that produce the brown/black pigment eumelanin (Garibyan et al, 2010; Schwarz and Schwarz, 2011). Interestingly, melanin production is ubiquitous across all kingdoms, and in insects, melanin synthesis is used for signalling (aposematism), camouflage and defence against pathogens (Whitten and Coates, 2017). A myriad of studies, including those with *Drosophila melanogaster* and *Galleria mellonella*, have demonstrated that melanisation is a key part of the insect innate immune system, and is deployed at the site of injury to help seal wounds and clear an infection from a host (Uçkan et al, 2010; Tang, 2009; Dubovskiy et al, 2013; Binggeli et al, 2014; Grizanova et al, 2019; Smith et al, 2022).

To understand the influence that UV radiation exposure has on the innate immune system, we used insect larvae from the Greater wax moth, *Galleria mellonella*. Wax moth larvae are ethically more acceptable than vertebrates for experimentation (i.e., rodents) and larger than traditional invertebrate models like Drosophilids and nematodes; requiring neither specialist equipment nor training (Mather, 2001; Mylonakis, 2008; Lionakis, 2011; Coates et al, 2019; Emery et al., 2019, 2021; Lim et al, 2022). *Galleria mellonella* possesses an innate immune system – lacking the ability to produce clonal immunoglobulins or long-term immune memory (adaptive immunity) – thereby permitting a targeted assessment of UV exposure on frontline immune defences. Like mammalian immunity, *G. mellonella* innate immunity consists of both cellular and humoral components, with the former comprising a heterogeneous population of circulating haemocytes like human and mouse neutrophils/macrophages (Browne et al, 2013). They play a critical role in pathogen phagocytosis and encapsulation (Kay et al, 2019; Krachler et al, 2021). The humoral defences comprise of soluble immune factors involved in activation of the pro-phenoloxidase cascade and have direct microbicidal and microbiostatic properties (Tsai et al 2016; Whitten and Coates, 2017).

Herein, our overall aim was to assess the impact of UV exposure on *G. mellonella* larvae using a series of whole organism, cellular and biochemical measures. We show that UV radiation affects cellular function in *G. mellonella* larvae, including circulating haemocyte numbers, melanisation levels, encapsulation rates, adhesion and metabolic activity. Under certain UV conditions, many larvae die and those that survive are more susceptible to bacterial (*Photorhabdus luminescens)* infection. Together, these data provide a baseline for investigating how other non-ionising radiation sources can influence the immune system in a suitable, non-vertebrate experimental system that can be used to study environmental stressors.

## MATERIALS AND METHODS

### *Galleria mellonella* maintenance

Final instar larvae of the greater wax moth, *G. mellonella*, were sourced from Livefoods Direct Ltd (UK) and stored in wood shavings (from the supplier) in the dark at 12°C. Healthy larvae weighing between 0.2 and 0.4 g and with little to no signs of melanisation were used in all experiments. Insect experimentation is not currently regulated in the UK, however, their use was approved by the University of Stirling’s Animal Welfare and Ethical Review Body (AWERB (16/17) 17 New Non-ASPA).

### UV irradiation of *Galleria mellonella*

Healthy larvae (average n = 27 per condition over 3 independent experiments) were irradiated with UV bulbs at either UVB (302 nm, XX-15MR Bench Lamp, UVP) or UVA (365 nm, XX-15LW Bench Lamp, UVP) at either a LOW (25 cm from bulb) or HIGH (10 cm) setting (Supplementary Figure 1). Larvae were exposed to UV for 60 and 120 minutes alongside control larvae not subjected to UV. Light intensity profiles for each position were determined using a RAMSES-ACC-UV/VIS-VA radiometer (TriOS, GmbH, Germany). (Supplementary Figure 2).

### Quantification of melanisation

After larvae were exposed to radiation, haemolymph was extracted through an incision behind the head using a hypodermic needle. Approximately 50 μl hemolymph per larva was collected and pooled in 0.5 ml ice-cold PBS (Fisher Scientific, 10388739, pH 7.4) and kept on ice to prevent spontaneous melanisation. Samples (100 μl) were transferred to a 96-well plate and absorbance at 405 nm was measured using a Molecular Devices VersaMax microplate reader with SoftMax Pro software (Kloezen et al, 2015). Seven to ten larvae per condition were used across two independent experiments.

### Total haemocyte counts

Ten microlitres of pooled, chilled haemolymph was mixed with an equal volume of PBS containing 0.4% (w/v) Trypan blue (MP Biomedicals), transferred to a haemocytometer (FastRead 102, Immune Systems) and examined under light microscopy conditions, and counted in triplicate.

### Haemocyte metabolic activity

Cell metabolic activity was assessed using the MTT [3-(4,5-dimethylthiazol-2-yl)-2,5-diphenyltetrazolium bromide] assay. Approximately, 1 × 10^5^ haemocytes from pooled haemolymph in Grace’s medium (G9771, Sigma-Aldrich, with 0.35 g/L NaHCO_3_, Antibiotic-Antimycotic, 0.1 mM phenylothiourea) were placed in triplicates in a 96-well plate and left for 30 min at room temperature to allow the cells to settle at the bottom of the well. Adhered cells were confirmed with an inverted microscope (XDS-2, Optika, Italy). The medium was replaced with 0.5 mg/ml MTT (15214654, Fisher Scientific) dissolved in PBS and left at room temperature for 30 min. Viable cells possess active mitochondrial dehydrogenase that cleaves the tetrazolium rings of the pale yellow MTT to form dark blue formazan crystals. Haemocytes were solubilised with dimethyl sulfoxide and read using a microplate reader set at a wavelength of 595 nm (VersaMax, Molecular Devices). The number of viable cells was directly proportional to the level of the formazan product created. Haemolymph of larvae that were not treated with UV were used for the positive control (for 100% viability). For the negative control, haemolymph was treated with 0.1% Triton-X solution before the addition of MTT (for 0% viability; as Lim et al, 2012).

### Haemocyte adhesion assay

Conditions for the adhesion assay were adapted from Humphries (2009). Approximately, 1 × 10^5^ haemocytes from pooled haemolymph were seeded into a 96-well plate in triplicates and left to settle at the bottom for 20 min at room temperature. PBS was removed and any unattached cells were washed once with PBS. Cells were fixed with 2% (w/v) glutaraldehyde for 10 min, washed once with PBS, stained with 0.1% (w/v) crystal violet, solubilized with 10% (v/v) acetic acid, and the absorbance was read at 570 nm (VersaMax, Molecular Devices). Positive and negative controls included cells from non-UV radiated larvae and cells pre-treated for 30 min with 5 μM cytochalasin D, respectively.

### Growth and maintenance of *Photorhabdus luminescens* subsp. *laumondii TT01*

*Photorhabdus luminescens* subsp. *laumondii* TT01 was obtained from NCIMB (#14339) and single colonies were cultured in Lysogeny Broth (LB) at 30°C in a rotational incubator (SSL4, Stuart) shaking at 150-220 rpm for 16 h prior to further processing. When required,1 ml of bacterial suspension was washed in PBS and the optical density (OD) was recorded (WPS, Biochrom) at 600 nm. Samples were diluted to OD values of 0.0001 and 0.00001 in PBS, which represents ∼4.5 ×10^6^ – 4.5 ×10^5^ colony forming units (CFUs) per ml prior to experimental use.

### *Galleria mellonella* infection with *Photorhabdus luminescens* TT01

Irradiated and non-irradiated larvae were injected into the last left proleg with 20 μl of bacterial cultures at OD 0.0001 (4.5 ×10^6^ CFU/ml) or 0.00001 (4.5 ×10^5^ CFU/ml) using a 1 ml syringe with a 21G hypodermic needle, representing 9.0 ×10^4^ or 9.0 ×10^3^ CFU/larva, respectively. At least 30 larvae were injected per condition. Larvae were incubated at 30 °C.

At 24 h post-injection, larvae were imaged using the ChemiDoc imaging system (Bio-Rad), with 10 s exposure time for all samples. Intensities of luminescence were determined by densitometry using ImageJ by freehand drawing around each larva and measuring the intensities therein. Bars are means ± SE from 15-18 larvae/condition. Statistical significance compared to non-irradiated control larvae (also challenged with bacteria) was determined by two-way ANOVA, and Dunnett’s multiple comparisons test (****, p ≤ 0.0001; ***, p ≤ 0.001; **, p ≤ 0.01; ns, not significant).

### Encapsulation assay

Five dishes containing healthy larvae were irradiated at either UVB (302 nm) or UVA (365 nm) at LOW (25 cm from bulb) or HIGH (10 cm) settings, for 60 or 120 min alongside control larvae not subjected to UV. Next, larvae were surface sterilised with 70% ethanol and a 3 mm piece of nylon thread (0.45 mm diameter, 100% polyamide, efco creative GmbH) was inserted gently into the haemocoelic cavity parallel to the GI tract via its last right pro-leg using a pair of fine forceps. Larvae were incubated for 24 h at 30°C, frozen at -20°C to kill the insect, defrosted and dissected to retrieve the nylon implant. Implants were imaged – at both ends – using an upright (light) microscope (CX31, Olympus) with an eyepiece camera (BF960, Swift Optical Instruments Ltd). Percentage of nylon thread covered by capsule formation was determined by measuring capsular area using ImageJ software (National Institutes of Health) and related to the entire thread as visualised (Lim et al, 2022). Sample sizes included 6 larvae per condition across two independent experiments.

## RESULTS

### Impact of UV radiation on *Galleria mellonella* health

Survival was not impacted negatively by subjecting larvae to either dose (low, high) of UVA for up to two hours, and was indistinguishable from the control group; 100% survival in each case after seven days. All larvae treated with the high dose of UVB (for two hours) died after 72 h (Supplementary Figure 3) – whereas those exposed to the low dose of UVB all survived. Next, we determined whether UVA or UVB enhanced insects’ susceptibility to infection by injecting post-irradiated larvae with *Photorhabdus luminescens*, a known bacterial pathogen of *G. mellonella* (e.g., Hu and Webster, 2000). At 16 h post-infection, larvae were imaged for luminescence produced by the bacterium, which gives an indication of the extent of septicaemia within the larval haemocoel (body cavity; Figure 1). Levels of bacterial dissemination varied across the larvae with average grey values higher than the uninfected control group. The highest mean grey value was from larvae exposed to UVB at the low dose prior to infection (Figure 1). Overall, both UVA (*X*^2^_(5)_ = 159.5, P < 0.0001) and UVB (*X*^2^_(5)_ = 183, P < 0.0001) exposure significantly increased *G. mellonella’s* susceptibility to infection (Figure 2) – UV exposed insects succumbed to bacterial infection faster than those not exposed to UV (regardless of the UV dose or duration). Median survival time for non-irradiated, infected insects was 72 hours, whereas irradiated insects were likely to die within 48 to 60 hours for UVA, and 48 hours for UVB (Supplementary Table 1). By 24 h post-infection, about a quarter to a third of UV irradiated larvae died (Figure 2). This trend continued for another 24 h before all larvae from all treatment groups died at 72 h. Duration of exposure to UV appears to be the proximal driver in enhancing insect susceptibility to infection; survival levels of *G. mellonella* exposed to UVA and UVB for 120 minutes, regardless of the dose, were significantly different to the positive control group and 60-minute exposure groups over the experimental period (see Table 1 for all comparisons).

**Table 1.**
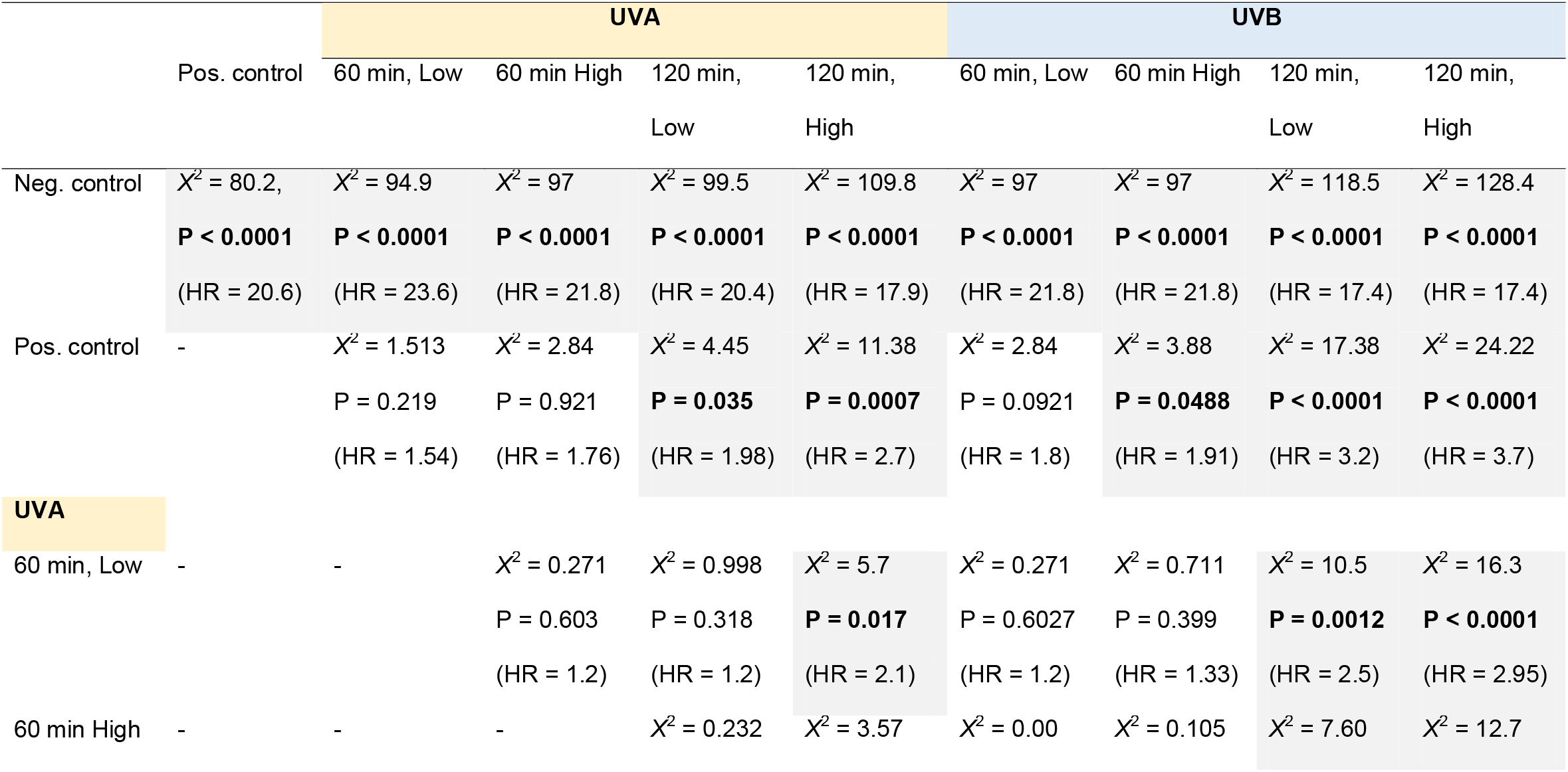
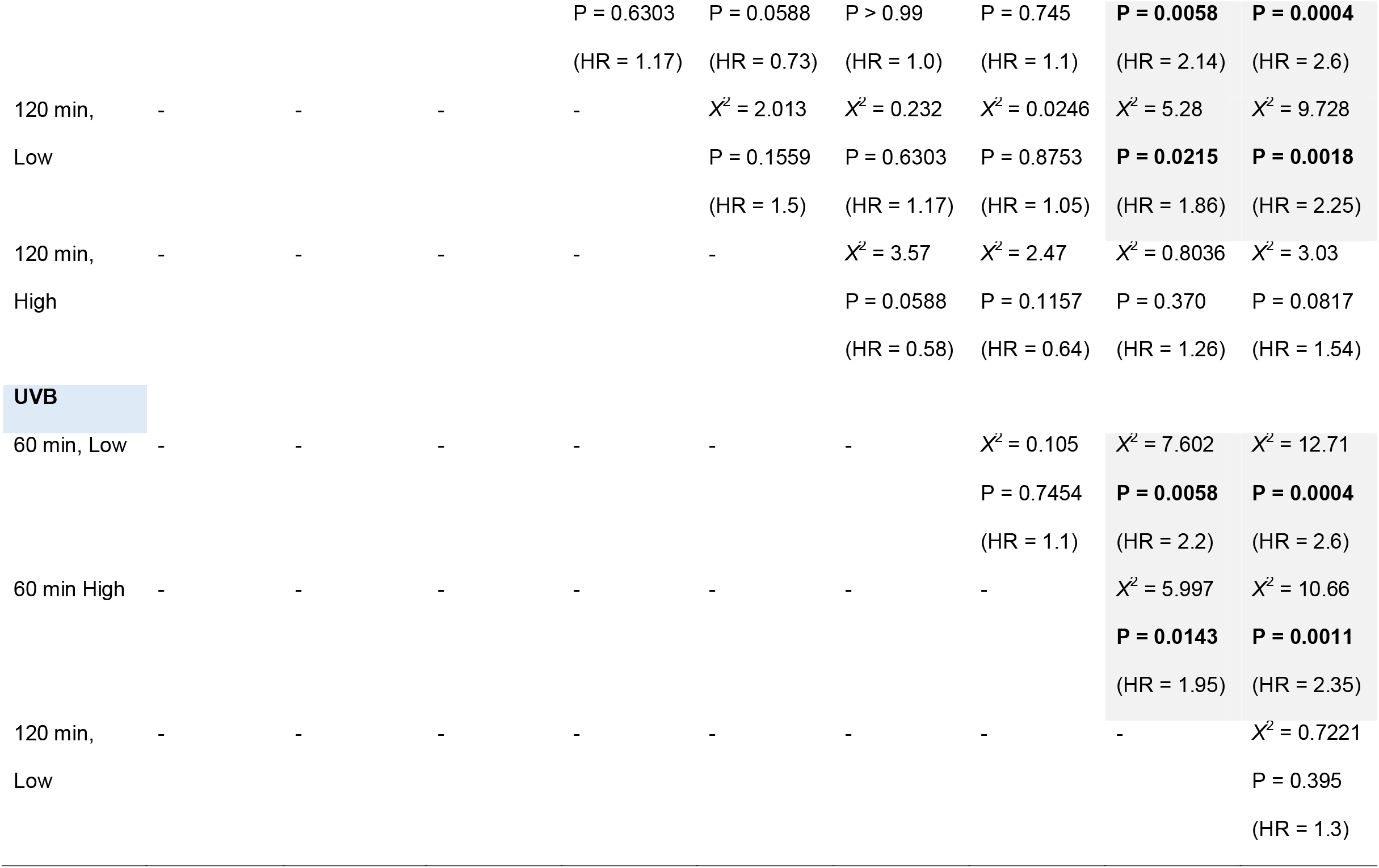
Survival analyses of *G. mellonella* larvae pre-exposed to UV radiation (A and B) for either one or two hours, then infected 24 hours later with *P. luminescens*. Log rank (Mantel Cox) tests were used to compare curves (df = 1 in all cases). Hazard ratios (Mantel Haenszel) are presented in parentheses. Negative control = non-radiated, **uninfected** larvae. Positive control = non-radiated, **infected** larvae. UVA = 365 nm. UVB = 302 nm.

**Figure 1.**
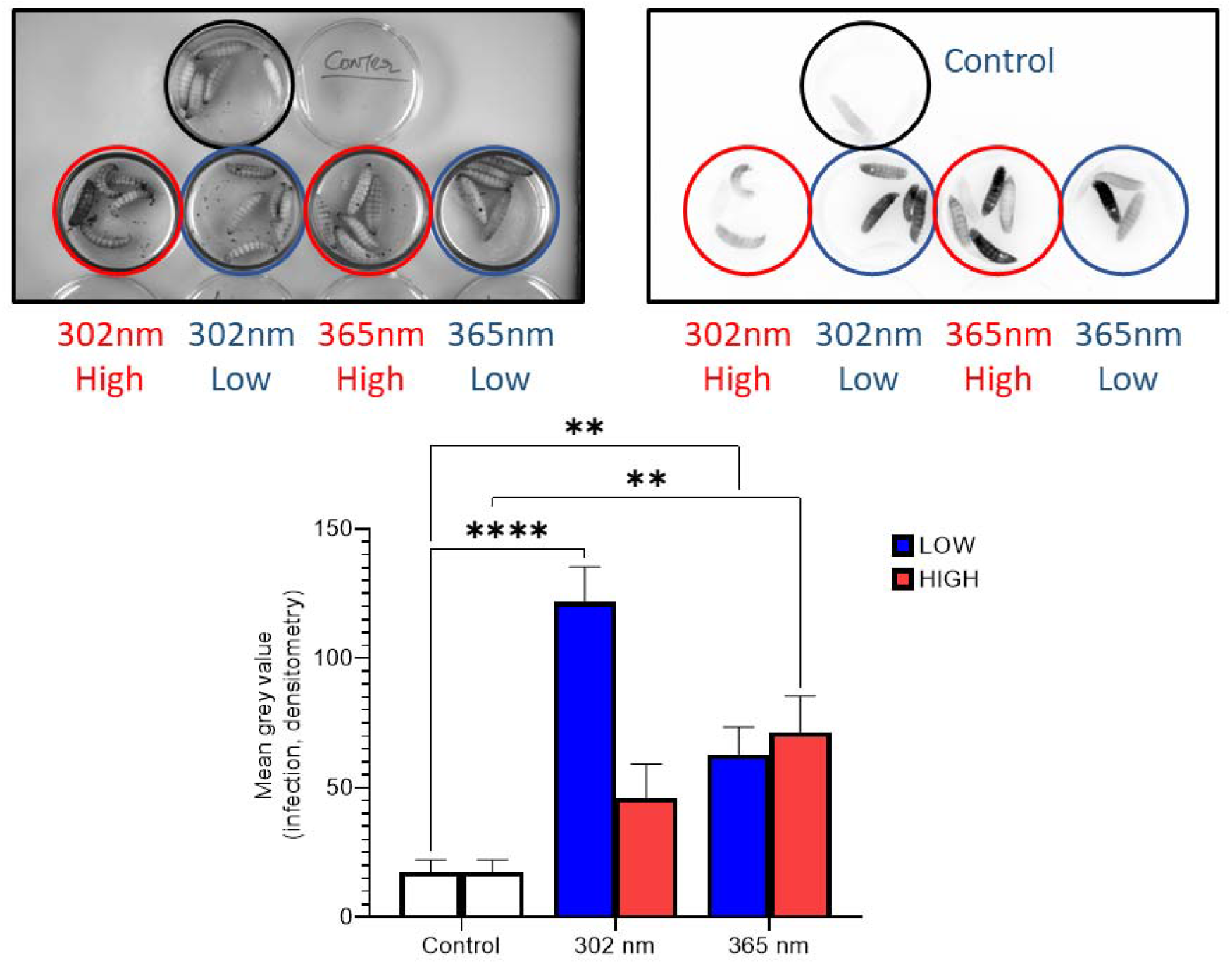
Proliferation of *Photorhabdus luminescens* in UV irradiated *Galleria mellonella*. Larvae were exposed to UVA (365 nm) and UVB (302 nm) at two different settings for 60 min, before a challenge with *P. luminescens* TT01. After 24 hr, larvae were imaged, and intensities of luminescence were determined by densitometry using ImageJ as described in the Materials and Methods section. Bars are means ± SEM from 15-18 larvae/condition. Statistical significance compared to non-irradiated control larvae (also challenged with bacteria) was determined by two-way ANOVA, and Dunnett’s multiple comparisons test. (****) p ≤ 0.0001, (**) p ≤ 0.01.

**Figure 2.**
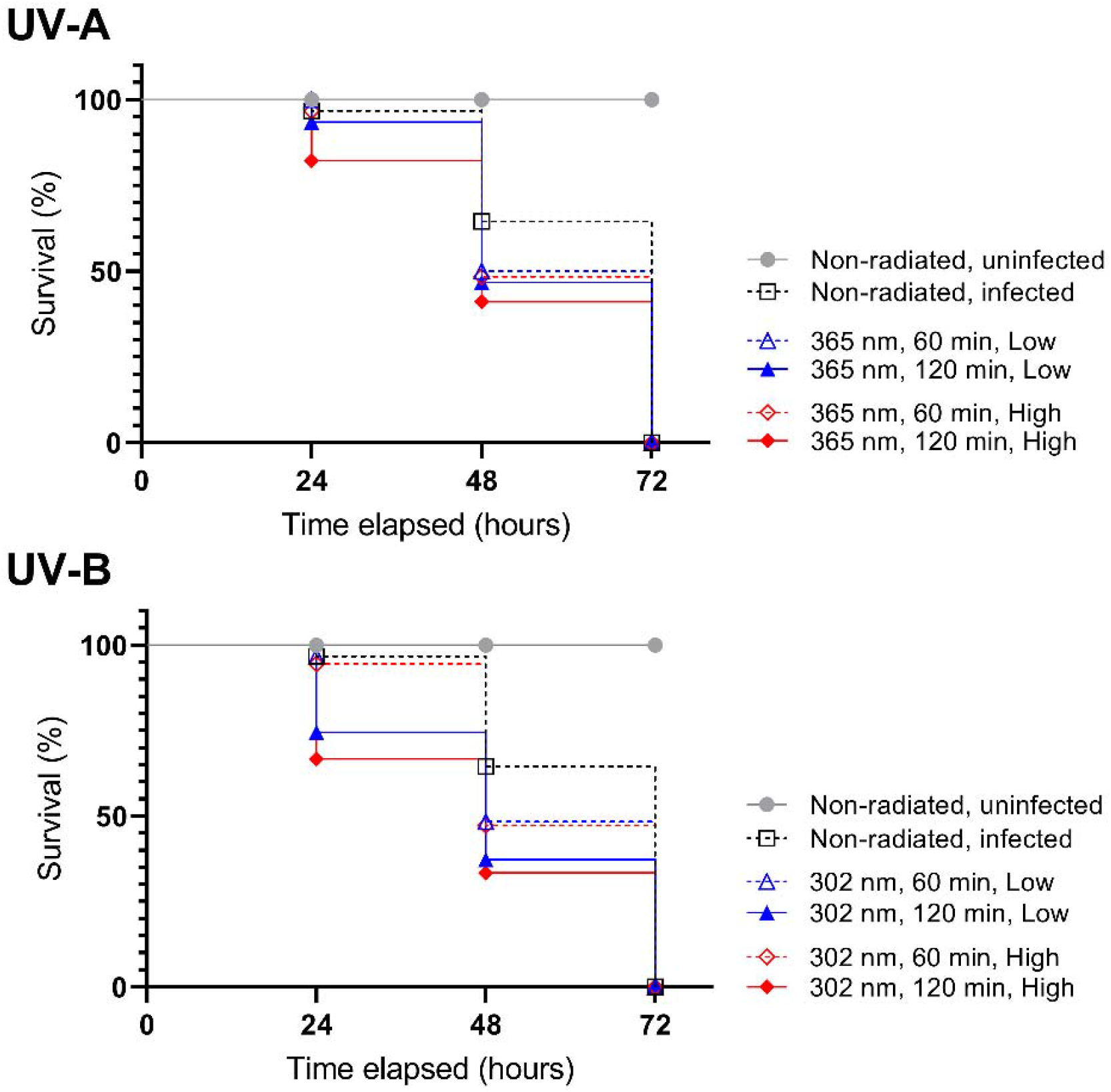
Exposure to UV radiation increases *Galleria mellonella* susceptibility to *Photorhabdus luminescens* infection. Thirty larvae per group were exposed to UV (A and B) radiation (365 nm and 302 nm, respectively) for 60 or 120 minutes at two doses (low versus high) before being injected with 20 μl of either PBS (negative control), or a suspension of PBS containing 9.0 ×10^4^ or 9.0 ×10^3^ CFU/larva of *P. luminescens* WT strain. Larvae were monitored every 24 h for 3 days. Data are means ± SE.

### Impact of UV radiation on *Galleria mellonella* gut and faecal content

Previously, larvae exposed to Cs-137 ionising radiation led to increased faecal load over time (Lim et al., 2022). To determine what effect non-ionising radiation had on the insect gut and its bacterial flora, larvae were irradiated, incubated for 3 days and were monitored for changes in faecal discharge. There was a significant decrease in the amount of faeces produced by larvae exposed to UVB for 120 min at the high dose when compared to the control (0.28 ± 0.06g vs 1.01 ± 0.25g, *p* = 0.03; Figure 3A). Interestingly, we found significantly higher numbers of viable bacterial CFUs from the faeces produced by those UVB-exposed larvae compared to the control (Figure 3B; 8.7 ± 3.7 × 10^6^ vs 5.2 ± 1.8 × 10^6^, *p* = 0.001) with a number of samples having fewer viable bacteria compared to the control (UVB, 120 min, LOW: 1.6 ± 0.6 × 10^6^; UVA, 120 min, LOW: 1.9 ± 0.6 × 10^6^; UVA, 120 min, HIGH: 0.98 ± 0.3 × 10^6^, *p* > 0.05). This suggests UVB radiation has an indirect effect on the *G. mellonella* gut and its flora.

**Figure 3:**
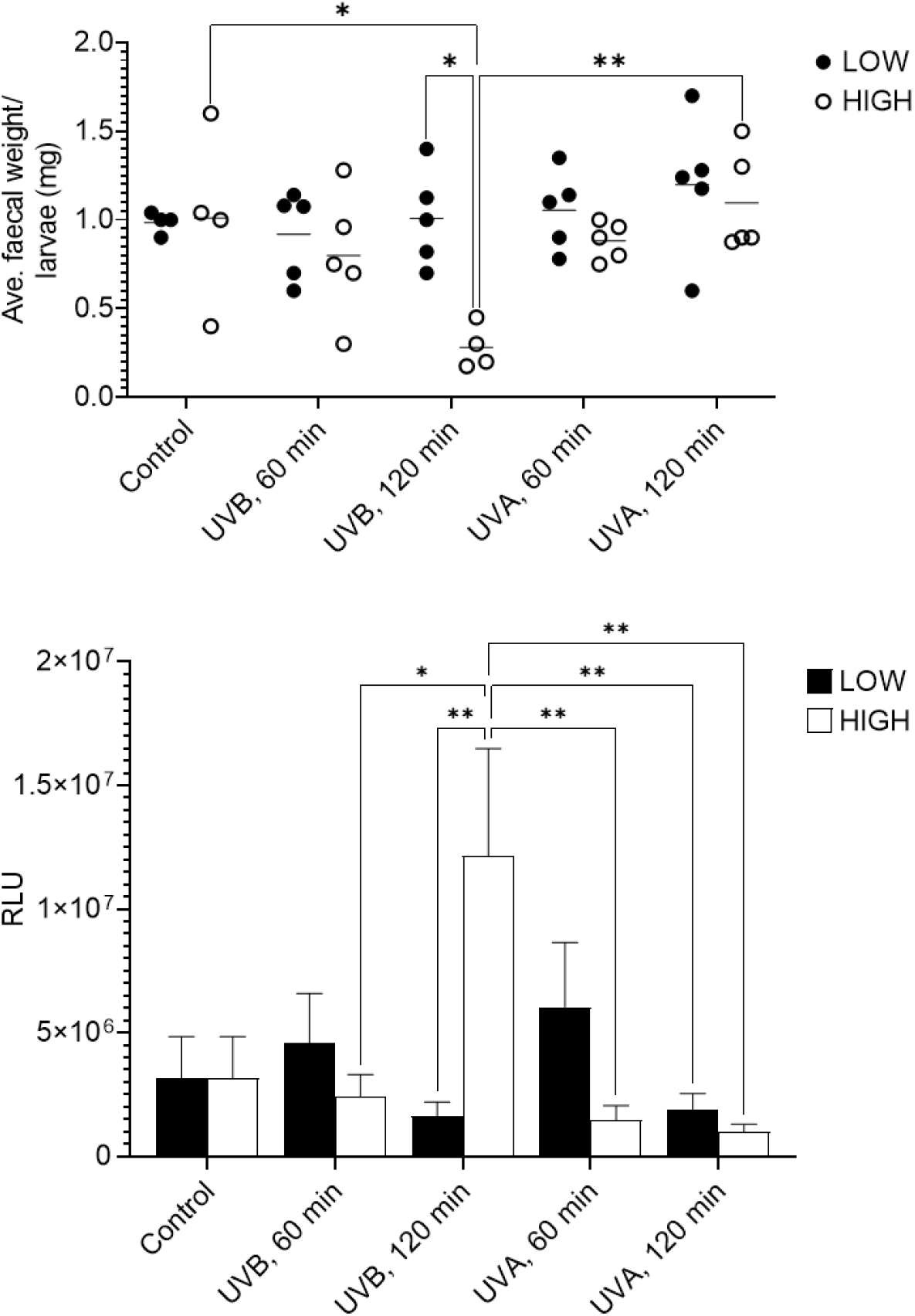
UV radiation affects faecal discharge from *Galleria mellonella*. Larvae were exposed to UVA (365 nm) or UVB (302 nm) radiation from UV lamps for the stated amount of time. Larvae were incubated and for 3 days and faecal material was weighed (**A**), resuspended with water and tested for viable bacteria (**B**), as described in the Materials and Methods. Significance was determined by two-way ANOVA, and a Tukey’s multiple comparisons test. (**) p ≤ 0.01, (*) p ≤ 0.05. Results are expressed as the mean ± SEM of at least three independent experiments (n = 5-7 per sample).

### UV radiation induces melanisation in insects

Broadly, the pigment melanin is produced in response to UV as it plays a role as a photoprotective factor in humans, resulting in darkening of the of skin in the short term or promoting cancer in the long term (Brenner and Hearing, 2008). Conversely, melanogenesis in insects has different functions, notably in defence against pathogens and parasites. Regardless of the dose (low, high), UV source (A, B), or duration of exposure (60 and 120 minutes), we recorded elevated levels (≥0.2 A405 nm) of melanotic (quinone) content in the haemolymph of treated *G. mellonella* compared to the negative control (≤0.18 A405 nm; Figure 4). Significant increases in melanin were recorded in larvae exposed to UVB for 120 min at both Low (0.26 ± 0.02; *p* = 0.03) and High (0.28 ± 0.02; *p* = 0.0009) settings, when compared to the unexposed control (Figure 4).

**Figure 4:**
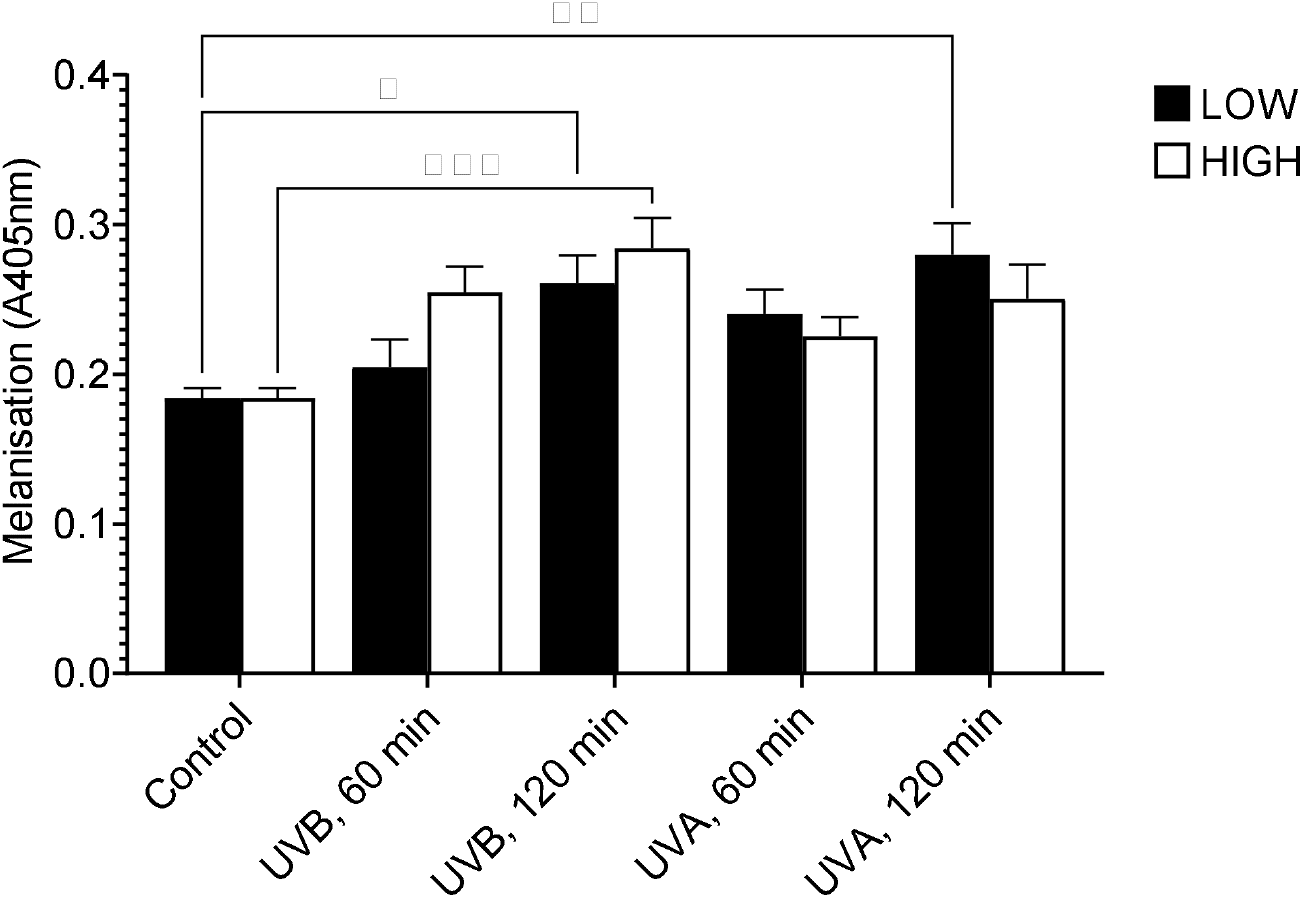
UV radiation promotes melanisation. Larvae were exposed to UVA (365 nm) or UVB (302 nm) for either one or two hours priort to haemolymph extraction.Melanin precurors, ie. dopachrome signals were recorded at 405 nm. Bars are means ± SE from three independent experiments (n = 20). Statistical significance was determined by two-way ANOVA, and Tukey’s multiple comparisons test. (****) p≤ 0.0001, (***) p≤0.001, (*) p ≤ 0.05.

### Impact of UV radiation on haemocytes numbers, viability and activity

Fluctuating haemocyte numbers in the haemolymph is an indication of the immune response of *G. mellonella* (Mowlds et al. 2010). We counted freely circulating haemocytes from larvae subjected to UV radiation and found those who were exposed to UVA for 60 min at Low and High doses yielded the highest cell counts (21.7 ± 3.1 × 10^5^ and 31.2 ± 8.4 × 10^5^, respectively). After 120 minutes, haemocyte numbers decreased significantly (p = 0.028) to below control levels for the high dose but remained stable for the low UVA dose. With UVB, total haemocyte counts showed little differences or were lower than the controls, though not significantly so (15.7 ± 1.0 × 10^5^ versus 9.3 ± 3.2 × 10^5^; p = 0.9; Figure 5A). Circulating haemocytes in the haemolymph of larvae showed differential responses to UV wavelength and exposure.

**Figure 5.**
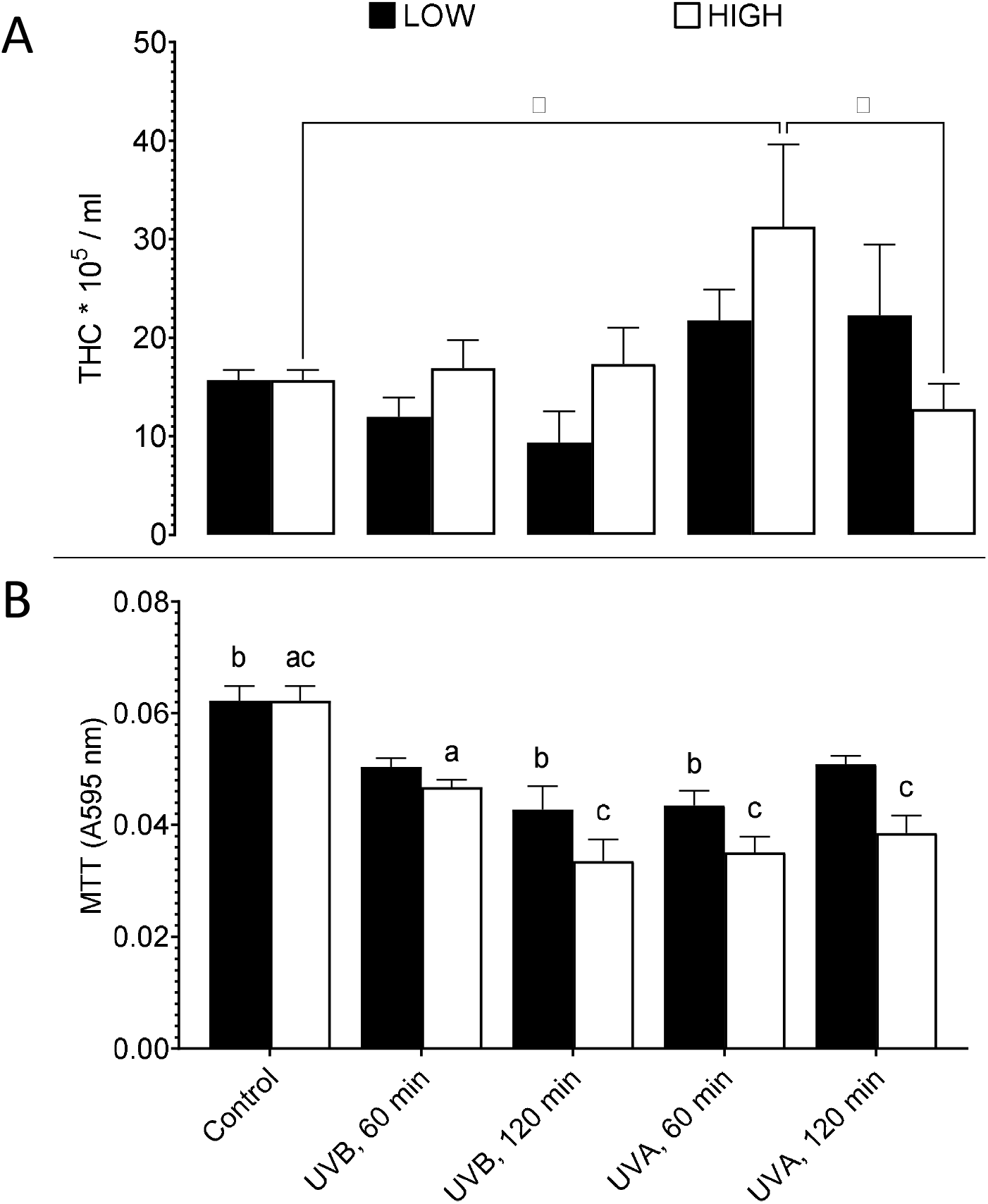
UV radiation affects total hemocyte counts and metabolic activity in *Galleria mellonella* haemocytes. Larvae were exposed to UVA (365 nm) or UVB (302 nm) for the stated time before hemolymph was extracted, pooled into PBS and counted by trypan blue exclusion with a hemocytometer (**A**) or measured for metabolic activity spectrophotometrically (**B**) as described in the Materials and Methods section. Bars are means ± SEM from 6-13 larvae/condition,. Statistical significance was determined by two-way ANOVA, and Tukey’s multiple comparisons test (*, p ≤ 0.05) (**A**) or shared letters represent significant differences (a) p ≤ 0.02; (b) p ≤ 0.0005; (c) p ≤ 0.0001 (**B**). Results are expressed as the mean ± SEM of 2 – 3 independent experiments (n = 7 – 9).

Across all irradiated samples, there was a broad, significant decrease in MTT levels when compared to the non-irradiated control (Figure 5B) - a proxy for cell viability (Kamiloglu et al, 2020). Haemocytes from larvae exposed to UVB showed both time- and dose-dependent decreases, with the lowest MTT value of ∼0.03 recorded after 120 minutes at the high dose, compared to the respective control value >0.06. This suggests that the higher cell counts could be made up of a large population of cells containing compromised (or downregulated) mitochondrial function (Figure 5).

Cell adhesion is a critical function that occurs during cell-cell interactions or cell-extracellular matrix interactions via cell-adhesion molecules, in response to changes in its surroundings. In humans, changes to cell adhesion can disrupt cellular functions and lead to a variety of diseases, including cancer, arthritis and infections (Harjunpää et al, 2019; Janiszewska et al, 2020; Johansson et al, 1999). *Galleria mellonella* larvae were exposed to UV radiation prior to haemolymph extraction and testing haemocyte adhesion to plastic using a crystal violet stain. As a positive control, haemocytes were treated with 5 mM cytochalasin D, an inhibitor of actin polymerisation, a process critical for adhesion (Banville et al., 2011*). Ex vivo* haemocytes from UVB-exposed insects showed significantly reduced adherent properties than control (unexposed) larvae (0.08 ± 0.02 and 0.21 ± 0.02, respectively; Figure 6A). When haemocytes from both irradiated and control insects were treated with cytochalasin D for 20 min, adhesion levels reduced to ≤0.05 ± 0.01 A570 nm (Figure 6A). There was a general decrease in adhesion to plastic with larvae that were UV-treated in dose- and time-dependent manners (*p* < 0.05 in all cases; Figure 6B) – most prominently observed with larvae exposed to UVA, at high setting for 60 min (0.10 ± 0.01, *p* = 0.003) and 120 min (0.11 ± 0.01, *p* = 0.01) when compared to the low setting (0.19 ± 0.01 and 0.19 ± 0.01, respectively) (Figure 6B). These data confirm that adhesion is an actin-dependent process for *G. mellonella* haemocytes, and that acute UV radiation compromises this cellular process.

**Figure 6:**
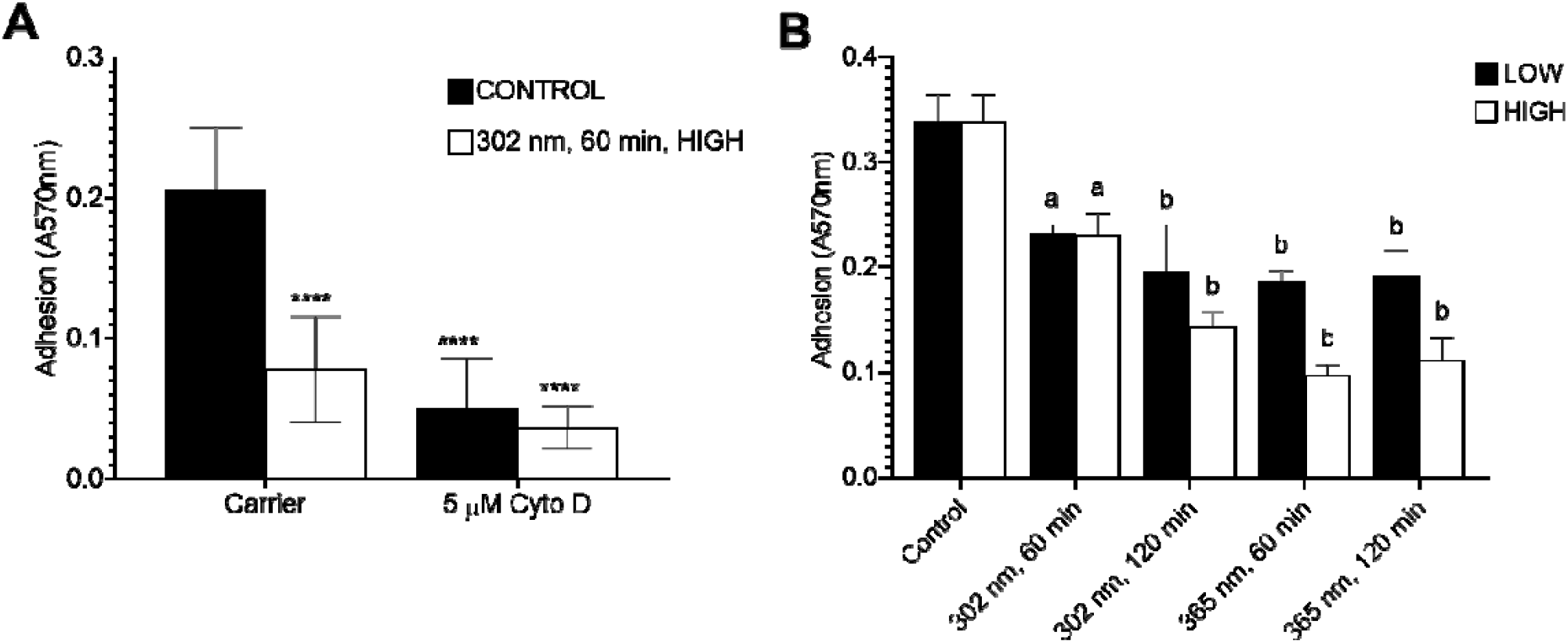
UV radiation affects haemocyte adhesion. Larvae were UV-irradiated, before haemolymph extracted, pooled and adhered to plastic, in multiple wells within a 96-well plate (A,B). As control, haemolymph was treated with 5 mM cytochalasin D for 20 min before adhering to plastic (A). Attached hemocytes were fixed with glutaraldehyde, stained with crystal violet and absorbance measured at 570 nm using a plate reader as described in the Materials and Methods section (A,B). Bars were means ± s.e.m. from 15-18 larvae/condition, after subtracting the values from the blank wells, across 3 independent experiments. Statistical significance was determined by two-way ANOVA, and Tukey’s multiple comparisons test. (****) p≤ 0.0001.

Additionally, *G. mellonella* larvae were UV-irradiated before inserting a piece of nylon thread into the haemocoel (body cavity) to test for responsiveness to non-self. UVB exposure coincided with decreased levels of haemocyte-mediated encapsulation – significant reductions for larvae treated with the high dose of UVB (60 min and 120 min: 3.98 ±1.2 and 5.7 ± 2.7, respectively; *p* < 0.05; Figure 7). Overall, UV radiation downregulates cell activity but has varying impact on cell numbers.

**Figure 7.**
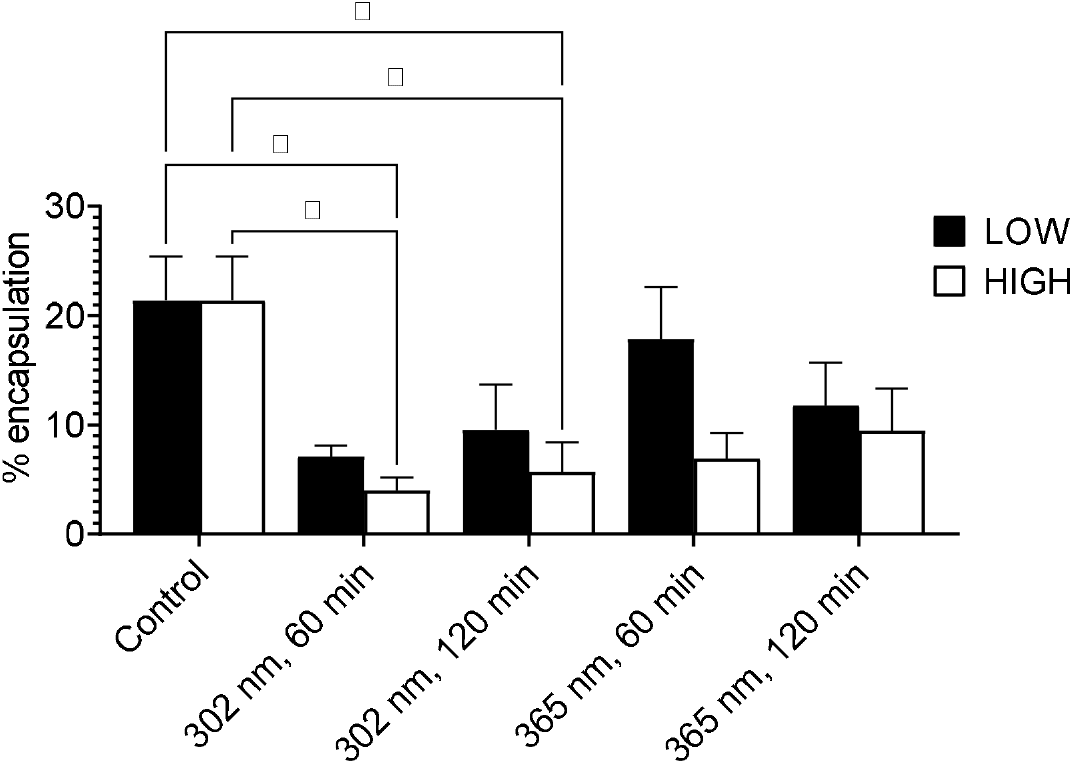
UV radiation affects encapsulation in *Galleria mellonella. G. mellonella* larvae were UV-irradiated for the stated time and wavelength, before inserting a piece of nylon thread into the haemocoelic cavity. Larvae were incubated at 25 °C for 24 hr, frozen, dissected and nylon thread removed and imaged under light microscopy as described in the Materials and Methods. % encapsulation was the surface area of thread covered by capsule material. Data were means ± SEM. Significance compared to irradiated larvae with non-radiated larvae was determined by two-way ANOVA, and Tukey’s multiple comparisons test. (*) p ≤ 0.05, otherwise not significant.

## DISCUSSION

The main source of non-ionising ultraviolet radiation is the sun and while the shortest range of UV wavelengths, UVC (< 280 nm), is absorbed by ozone in the stratosphere, UVB and UVA are not. Many studies exist covering the pleiotropic, dose-dependent effects of UV on mammalian immunity at both local and systemic levels and on chromophores and skin signalling pathways (Norval et al, 2008; Hart and Norval, 2021). However, their impact on invertebrates remains underexplored.

In this study, all larvae exposed to UVB (302 nm) radiation for two hours died. Larvae subjected to UV radiation also had variable haemocyte counts with UVA yielding the highest total haemocyte counts and UVB radiation leading to haemocyte counts that were lower than control larvae. While this may give an apparent contradiction of the subsequent MTT data which showed a general and significant decrease in MTT responses across all irradiated samples when compared to the non-irradiated control, these are two different experiments with the former based on cell counts, the latter on mitochondrial function. Therefore, the higher cell counts could be made up of a large population of cells containing downregulated mitochondria function. Interestingly, and consistent with our data, enhanced apoptosis in peripheral blood mononuclear cells (PBMCs) was observed in bodies exposed to UVA over UVB, possibly through increased pro-apoptotic protein production due to photosensitised oxygen radicals (Narbutt et al, 2009; Hart and Norval, 2021). Furthermore, in *Galleria mellonella*, melanin production increased in the presence of UVA and UVB and yet its deposition along with other capsular material onto nylon thread is decreased compared to its non-irradiated controls. This is consistent with the damselfly (*Coenagrion puella*) model where a combination of UVA and UVB (mimicking summer) led to increased melanin deposition on the exoskeleton surface yet displayed reduced encapsulation of nylon filaments (Debecker et al, 2015). Melanin is a known UV photoadaptor and/or photoprotectant, and the elevated levels in insect haemolymph in response to UV is unsurprising. Separately, encapsulation is driven by haemocytes’ ability to adhere to, and ensheath a non-self (foreign) invader and we report haemocyte mediated encapsulation is compromised in UV-treated insects – likely due to fewer haemocytes present, and which are dysfunctional (low MTT levels, and incapable of adherence).

Finally, we found UV radiation decreases haemocyte adhesion to plastic *ex vivo*. Again, this is consistent with data from human studies which showed that UVB irradiation of the human body led to decreases in both phagocytic, chemotactic and <u>adhesion</u> activities of neutrophils, largely due to decreased expression of receptors involved in such processes (Lundin et al, 1990; Leino et al, 1999; Hart and Norval, 2021). This further establishes the synonymy of cellular innate immunity in *G. mellonella* and mammals, and that the former is a candidate model for reducing our reliance on mammals for such studies (Browne et al 2013; Kay et al 2019; Krachler et al, 2021; Lim et al, 2022). Therefore, while more haemocytes remain in larvae exposed to UVA, they were functionally compromised, whereas UVB-radiated larvae had fewer haemocytes which was also compromised – which UVB is more potent an antagonist – as reflected in larval health, and expedited mortality when infected, in addition to compromised haemocyte functionality.

Larvae exposed to UVB radiation had decreased faecal discharge, compared to UVA and controls. Yet, interestingly, the faecal droppings from that sample group had the highest levels of viable bacteria. UV has little penetrating power, so the impact on the gut flora is likely an indirect (i.e., knock on) effect. We can speculate that the gut contents are emptied due to shock (thereby accounting for the volume of faecal discharge), so the bacteria are still viable, whereas the bacteria are non-viable in the other treatments because the gut contents remain *in situ* post UV exposure. Previously, gut bacteria from several families were enriched with altered microbial profiles both in mammals and insects exposed to control and increasing doses of UVB or any environmental toxins such as okadaic acid (Ghaly et al, 2018; Bosman et al, 2019; Emery et al, 2021). This increase in faecal microbiota composition and diversity is both beneficial and detrimental to the larvae. This change was suggested to be driven by both innate and adaptive immune cells (Clark and Mach, 2016). These immune cells traffic to the gut, release mediators and change the composition and diversity of the gut microbiome (e.g., Marra et al, 2021). Interestingly, a series of signalling cascades occurs between insect and fungal molecules produced in battle during the binding and invading of entomopathogenic fungal conidia to an insect integument (Mukherjee and Vilcinskas, 2018). Moreover, *G. mellonella* produces antimicrobial peptides (AMP) that regulates gut microbiota and the higher levels of viable bacteria in the. The higher percentage of viable bacteria in the faeces may be due to impaired AMP synthesis upon UV-treatment (Dubovskiy et al, 2016). Therefore, our study, while using an insect model that lacks an adaptive immune system, complements those conclusions from mammalian and insect studies and suggests the existence of a novel integument-gut axis in insects.

Finally, the perturbation of *G. mellonella*’s immunity by UV irradiation meant that this insect was more susceptible to infection by allowing the proliferation of *Photorhabdus luminescens*, an entomopathogenic enterobacterium. UVB has a negative impact on frontline (haemocyte) immunity in insect larvae, and likely explains why the colonisation of the haemocoel by *P. luminescens* is so rapid and reduces the time to *in exitus* by 24 hours.

While the exact mechanism is not clearly understood, in mammals, there is a balance between immune response mounted by the host versus bacterial pathogenesis. Any changes to the activation of immune responses (e.g. by UV) can be counter-productive to the host, and thereby allowing greater pathogenesis. Hence, pathogenicity is a function of both microbial traits and host’s immune response. This integrated view of pathogenicity in insects termed the “damage threshold hypothesis” which is reminiscent of the “damage response framework” observed in mammals (Moreno-García et al, 2014; Casadevall and Pirofski, 2003; Krachler et al., 2021). We suggest that *G. mellonella* larvae are not only suitable as a tractable model organism to study cellular function when exposed to UV radiation, but also to better probe the full spectrum of microbe–host-environment interactions in insects. Together, these data provide opportunities for investigating how other environmental stressors can influence the immune system using this simple, but highly effective experimental system.

## ACKNOWLEDGEMENTS

JL would like to thank his family for their help and support and to Sabine Matallana-Surget for the use of her UV lamps. We acknowledge the University of Stirling and the Carnegie Trust for the Universities of Scotland (RIG008296) for some funds. CC would like to thank Susan and Lonán Coates for their support. Special thanks to Ryan Ferguson (Nuffield Foundation) and Matthew Keddie for gathering some preliminary data.

## SUPPLEMENTARY INFORMATION

**Supplementary Figure 1.**
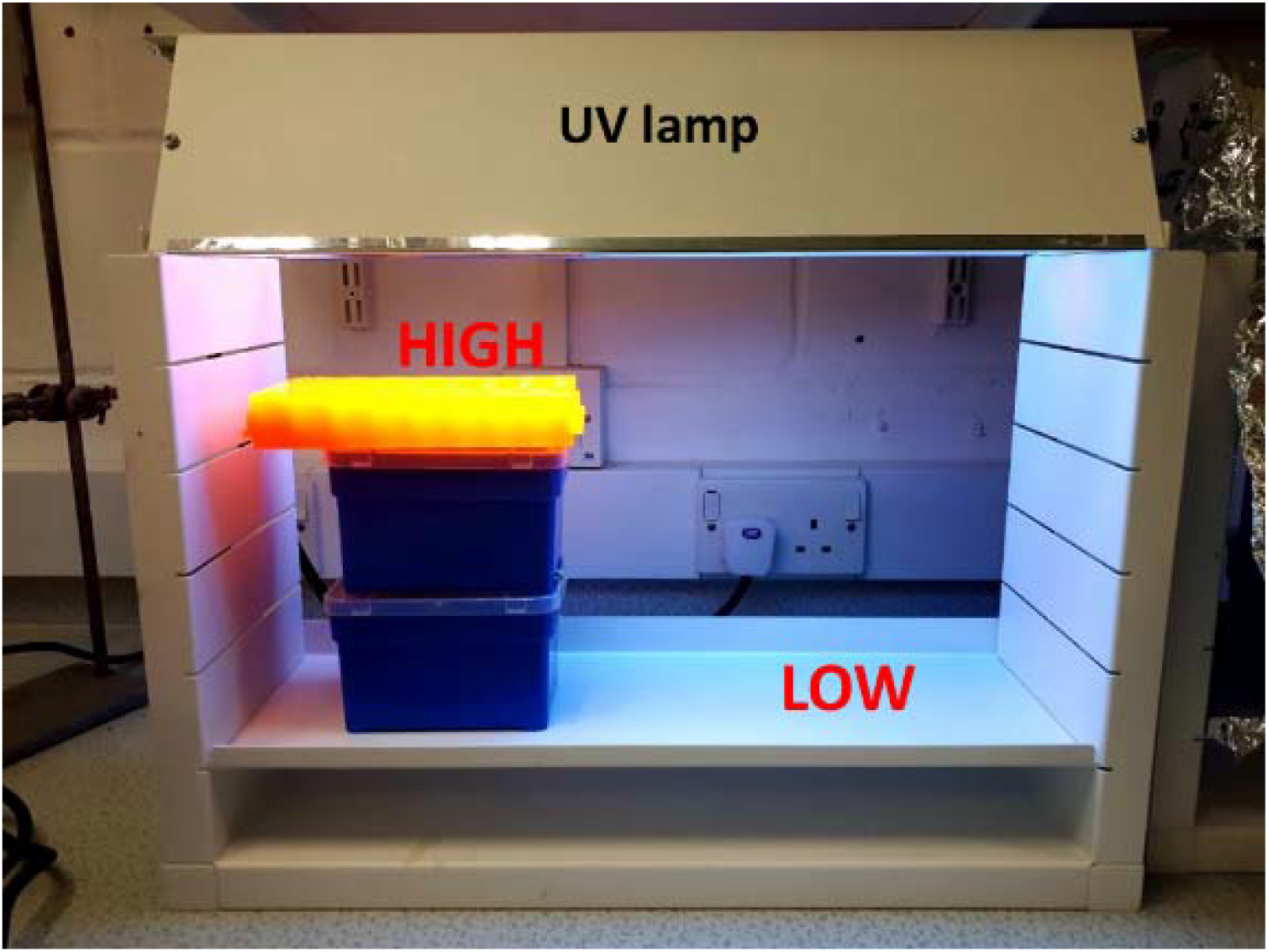
Experimental set up of *Galleria mellonella* larvae under UV lamps. Larvae exposed to UV lamps (either 302 or 365 nm) at either HIGH (10 cm from bulb) or LOW (25 cm) settings, depending on the distance from the source.

**Supplementary Figure 2.**
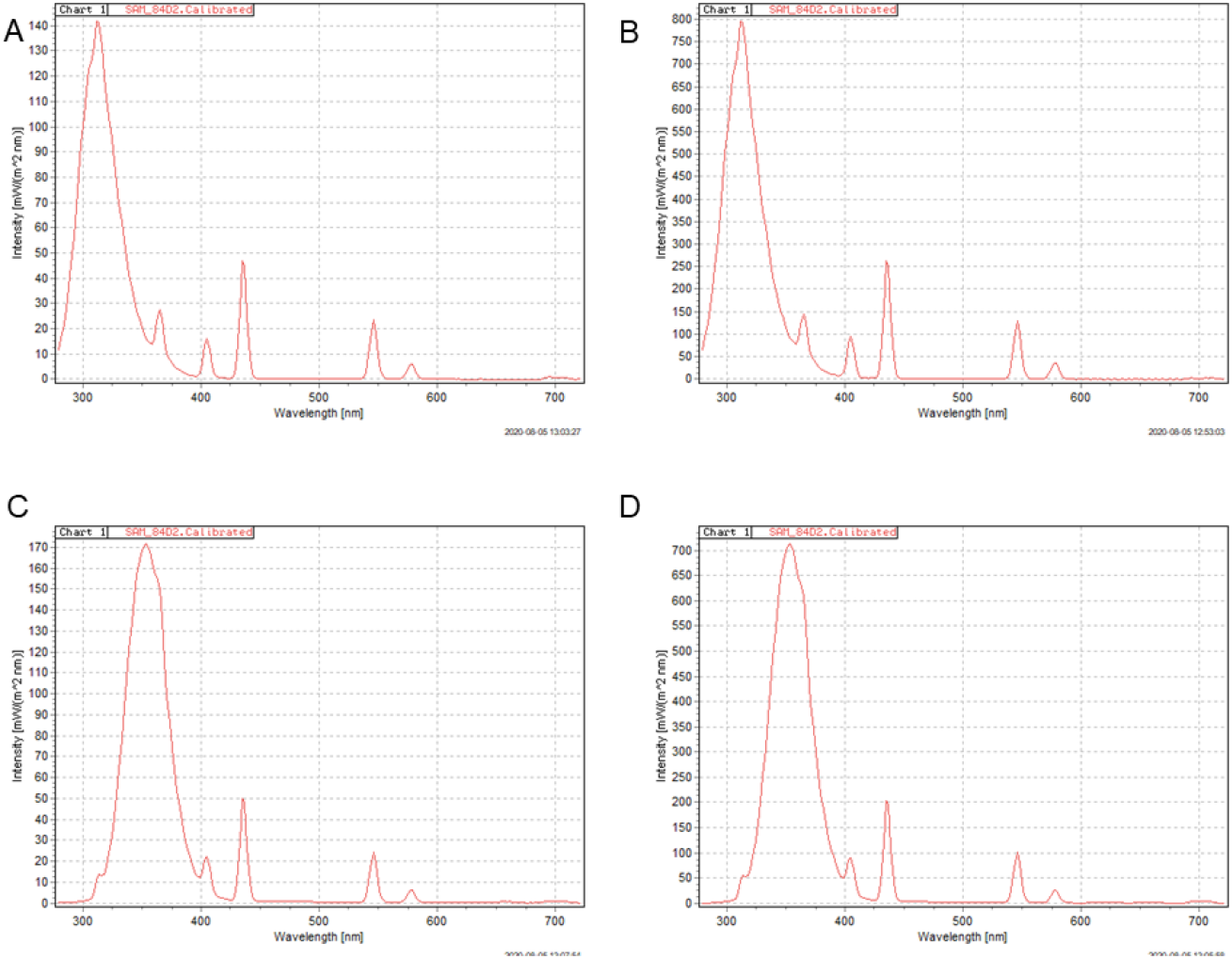
Light intensity profiles emitted from bulbs at different heights. UV light intensities experienced at 2 different positions (Low, A and C; High, B and D; see Figure 1) at 2 different lamp sources (302 nm, A and B; 365 nm, C and D) were measured across the light spectrum using a RAMSES-ACC-UV/VIS-VA radiometer (TriOS, GmbH, Germany).

**Supplementary Figure 3.**
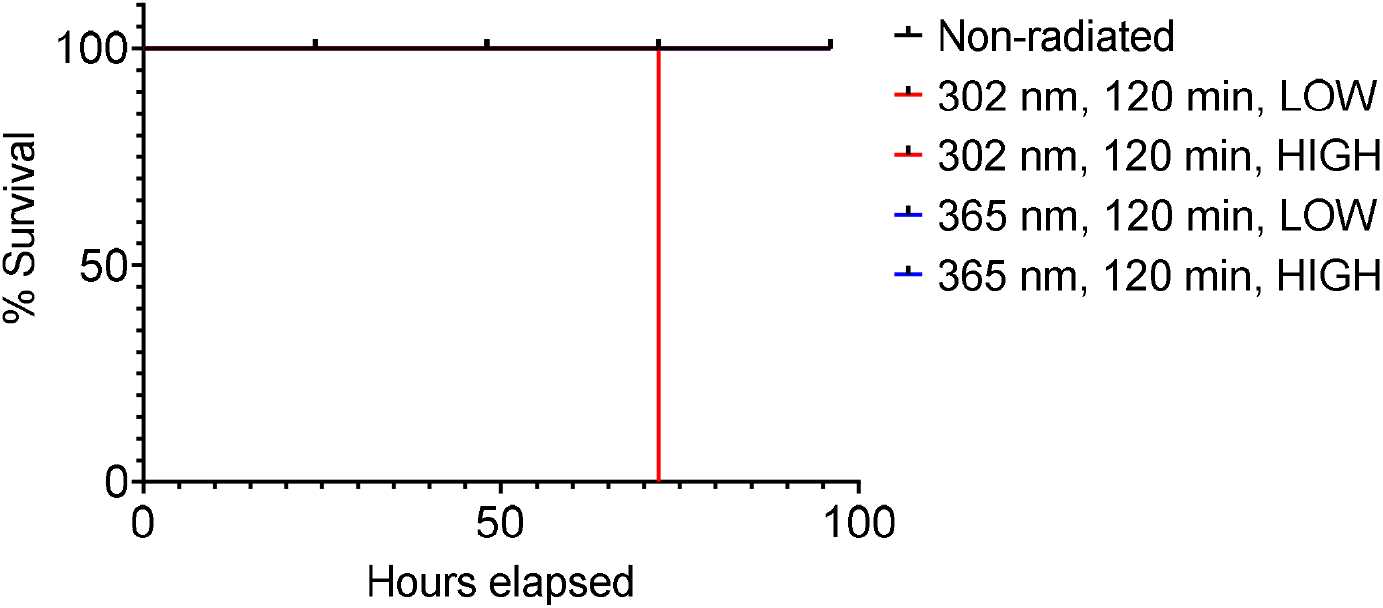
Impact of UV radiation on *Galleria mellonella* viability. Ten larvae were incubated at 25 °C for 24 h before exposure to UV radiation (302/365 nm) for 120 min and monitored every 24 h for 7 days for survivability.

**Supplementary Table 1.** Median survival times for UV-irradiated insect larvae infected with *P. luminescens* (i.e., probability of survival equalling 50%)

|  | Median survival time |
| --- | --- |
| **Controls** |  |
| Non-radiated, uninfected (negative control) | Not defined |
| Non-radiated, infected (positive control) | 72 hours |
| **Wavelength, duration, dose (low vs. high)** |  |
| **UV-A** |  |
| 365 nm, 60 min, Low | 60 hours |
| 365 nm, 120 min, Low | 48 hours |
| 365 nm, 60 min, **High** | 48 hours |
| 365 nm, 120 min, High | 48 hours |
| **UV-B** |  |
| 302 nm, 60 min, Low | 48 hours |
| 302 nm, 120 min, Low | 48 hours |
| 302 nm, 60 min, High | 48 hours |
| 302 nm, 120 min, High | 48 hours |

## REFERENCES

Banville, N., Fallon, J., McLoughlin, K. & Kavanagh, K. (2011) Disruption of haemocyte function by exposure to cytochalasin b or nocodazole increases the susceptibility of Galleria mellonella larvae to infection. Microbes Infect, 13, 1191–8.

Binggeli, O., Neyen, C., Poidevin, M., & Lemaitre, B. (2014). Prophenoloxidase activation is required for survival to microbial infections in Drosophila. PLoS Pathogens, 10(5), e1004067.

Bosman, E. S., Albert, A. Y., Lui, H., Dutz, J. P. & Vallance, B. A. (2019) Skin Exposure to Narrow Band Ultraviolet (UVB) Light Modulates the Human Intestinal Microbiome. Front Microbiol, 10, 2410.

Brenner, M. & Hearing, V. J. (2008) The protective role of melanin against UV damage in human skin. Photochem Photobiol, 84, 539–49.

Browne, N., Heelan, M. & Kavanagh, K. (2013) An analysis of the structural and functional similarities of insect hemocytes and mammalian phagocytes. Virulence, 4, 597–603.

Casadevall, A. & Pirofski, L. A. (2003) The damage-response framework of microbial pathogenesis. Nat Rev Microbiol, 1, 17–24.

Clark, A. & Mach, N. (2016) Role of Vitamin D in the Hygiene Hypothesis: The Interplay between Vitamin D, Vitamin D Receptors, Gut Microbiota, and Immune Response. Front Immunol, 7, 627.

Coates, C. J., Lim, J., Harman, K., Rowley, A. F., Griffiths, D. J., Emery, H. & Layton, W. (2019) The insect, Galleria mellonella, is a compatible model for evaluating the toxicology of okadaic acid. Cell Biol Toxicol, 35, 219–232.

Copplestone, D., Coates, C. J. & Lim, J. (2022) Immune-modulatory effects of low dose γ-radiation on wax moth (Galleria mellonella) larvae. bioRxiv, 2022.10.11.511741.

Debecker, S., Sommaruga, R., Maes, T. & Stoks, R. (2015) Larval UV exposure impairs adult immune function through a trade-off with larval investment in cuticular melanin. Functional Ecology, 29, 1292–1299.

D’Orazio, J., Jarrett, S., Amaro-Ortiz, A. & Scott, T. (2013) UV radiation and the skin. Int J Mol Sci, 14, 12222–48.

Dubovskiy, I. M., Whitten, M. M., Kryukov, V. Y., Yaroslavtseva, O. N., Grizanova, E. V., Greig, C., Mukherjee, K., Vilcinskas, A., Mitkovets, P. V., Glupov, V. V. & Butt, T. M. (2013) More than a colour change: insect melanism, disease resistance and fecundity. Proc Biol Sci, 280, 20130584.

Dubovskiy, I. M., Grizanova, E. V., Whitten, M. M., Mukherjee, K., Greig, C., Alikina, T., Kabilov, M., Vilcinskas, A., Glupov, V. V. & Butt, T. M. (2016) Immuno-physiological adaptations confer wax moth Galleria mellonella resistance to Bacillus thuringiensis. Virulence, 7, 860–870.

Emery, H., Johnston, R., Rowley, A. F. & Coates, C. J. (2019) Indomethacin-induced gut damage in a surrogate insect model, Galleria mellonella. Arch Toxicol, 93, 2347–2360.

Emery, H., Traves, W., Rowley, A. F. & Coates, C. J. (2021) The diarrhetic shellfish-poisoning toxin, okadaic acid, provokes gastropathy, dysbiosis and susceptibility to bacterial infection in a non-rodent bioassay, Galleria mellonella. Arch Toxicol, 95, 3361–3376.

Garibyan, L. & Fisher, D. E. (2010) How sunlight causes melanoma. Curr Oncol Rep, 12, 319–26.

Ghaly, S., Kaakoush, N. O., Lloyd, F., Gordon, L., Forest, C., Lawrance, I. C. & Hart, P. H. (2018) Ultraviolet Irradiation of Skin Alters the Faecal Microbiome Independently of Vitamin D in Mice. Nutrients, 10.

Gonzalez Maglio, D. H., Paz, M. L. & Leoni, J. (2016) Sunlight Effects on Immune System: Is There Something Else in addition to UV-Induced Immunosuppression? Biomed Res Int, 2016, 1934518.

Halliday, G. M. (2005) Inflammation, gene mutation and photoimmunosuppression in response to UVR-induced oxidative damage contributes to photocarcinogenesis. Mutat Res, 571, 107–20.

Grizanova, E. V., Coates, C. J., Dubovskiy, I. M., & Butt, T. M. (2019). Metarhizium brunneum infection dynamics differ at the cuticle interface of susceptible and tolerant morphs of Galleria mellonella. Virulence, 10, 999–1012.

Harjunpaa, H., Llort Asens, M., Guenther, C. & Fagerholm, S. C. (2019) Cell Adhesion Molecules and Their Roles and Regulation in the Immune and Tumor Microenvironment. Front Immunol, 10, 1078.

Hart, P. H., Gorman, S. & Finlay-Jones, J. J. (2011) Modulation of the immune system by UV radiation: more than just the effects of vitamin D? Nat Rev Immunol, 11, 584–96.

Hart, P. H. & Norval, M. (2021) More Than Effects in Skin: Ultraviolet Radiation-Induced Changes in Immune Cells in Human Blood. Front Immunol, 12, 694086.

Hu, K. & Webster, J. M. (2000) Antibiotic production in relation to bacterial growth and nematode development in Photorhabdus--Heterorhabditis infected Galleria mellonella larvae. FEMS Microbiol Lett, 189, 219–23.

Humphries, M. J. (2009) Cell adhesion assays. Methods Mol Biol, 522, 203–10.

Janiszewska, M., Primi, M. C. & Izard, T. (2020) Cell adhesion in cancer: Beyond the migration of single cells. J Biol Chem, 295, 2495–2505.

Johansson, M. W. (1999) Cell adhesion molecules in invertebrate immunity. Dev Comp Immunol, 23, 303–15.

Kay, S., Edwards, J., Brown, J. & Dixon, R. (2019) Galleria mellonella Infection Model Identifies Both High and Low Lethality of Clostridium perfringens Toxigenic Strains and Their Response to Antimicrobials. Front Microbiol, 10, 1281.

Kloezen, W., van Helvert-van Poppel, M., Fahal, A. H. & van de Sande, W. W. (2015) A Madurella mycetomatis Grain Model in Galleria mellonella Larvae. PLoS Negl Trop Dis, 9, e0003926.

Krachler, A. M., Sirisaengtaksin, N., Monteith, P., Paine, C. E. T., Coates, C. J. & Lim, J. (2021) Defective phagocyte association during infection of Galleria mellonella with Yersinia pseudotuberculosis is detrimental to both insect host and microbe. Virulence, 12, 638–653.

Leino, L., Saarinen, K., Kivisto, K., Koulu, L., Jansen, C. T. & Punnonen, K. (1999) Systemic suppression of human peripheral blood phagocytic leukocytes after whole-body UVB irradiation. J Leukoc Biol, 65, 573–82.

Lim, J., Clements, M. A. & Dobson, J. (2012) Delivery of short interfering ribonucleic acid-complexed magnetic nanoparticles in an oscillating field occurs via caveolae-mediated endocytosis. PLoS One, 7, e51350.

Lionakis, M. S. (2011) Drosophila and Galleria insect model hosts: new tools for the study of fungal virulence, pharmacology and immunology. Virulence, 2, 521–7.

Lundin, A., Michaelsson, G., Venge, P. & Berne, B. (1990) Effects of UVB treatment on neutrophil function in psoriatic patients and healthy subjects. Acta Derm Venereol, 70, 39–45.

Mather, J. A. (2001) Animal Suffering: An Invertebrate Perspective. Journal of Applied Animal Welfare Science, 4, 151–156.

Marra, A., Hanson, M. A., Kondo, S., Erkosar, B. & Lemaitre, B. (2021) Drosophila Antimicrobial Peptides and Lysozymes Regulate Gut Microbiota Composition and Abundance. mBio, 12, e0082421.

Moreno-Garcia, M., Conde, R., Bello-Bedoy, R. & Lanz-Mendoza, H. (2014) The damage threshold hypothesis and the immune strategies of insects. Infect Genet Evol, 24, 25–33.

Mowlds, P., Coates, C., Renwick, J. & Kavanagh, K. (2010) Dose-dependent cellular and humoral responses in Galleria mellonella larvae following beta-glucan inoculation. Microbes Infect, 12, 146–53.

Mukherjee, K. & Vilcinskas, A. (2018) The entomopathogenic fungus Metarhizium robertsii communicates with the insect host Galleria mellonella during infection. Virulence, 9, 402–413.

Mylonakis, E. (2008) Galleria mellonella and the study of fungal pathogenesis: making the case for another genetically tractable model host. Mycopathologia, 165, 1–3.

Narbutt, J., Cebula, B., Lesiak, A., Sysa-Jedrzejowska, A., Norval, M., Robak, T. & Smolewski, P. (2009) The effect of repeated exposures to low-dose UV radiation on the apoptosis of peripheral blood mononuclear cells. Arch Dermatol, 145, 133–8.

Norval, M., McLoone, P., Lesiak, A. & Narbutt, J. (2008) The effect of chronic ultraviolet radiation on the human immune system. Photochem Photobiol, 84, 19–28.

Schwarz, T. (2002) Photoimmunosuppression. Photodermatol Photoimmunol Photomed, 18, 141–5.

Schwarz, A., Navid, F., Sparwasser, T., Clausen, B. E. & Schwarz, T. (2011) In vivo reprogramming of UV radiation-induced regulatory T-cell migration to inhibit the elicitation of contact hypersensitivity. J Allergy Clin Immunol, 128, 826–33.

Schwarz, T. (2010) The dark and the sunny sides of UVR-induced immunosuppression: photoimmunology revisited. J Invest Dermatol, 130, 49–54.

Schwarz, T. & Schwarz, A. (2011) Molecular mechanisms of ultraviolet radiation-induced immunosuppression. Eur J Cell Biol, 90, 560–4.

Smith, D. F. Q., Dragotakes, Q., Kulkarni, M., Hardwick, J. M. & Casadevall, A. (2022) Melanization is an important antifungal defense mechanism in <em>Galleria mellonella</em> hosts. bioRxiv, 2022.04.02.486843.

Sreevidya, C. S., Fukunaga, A., Khaskhely, N. M., Masaki, T., Ono, R., Nishigori, C. & Ullrich, S. E. (2010) Agents that reverse UV-Induced immune suppression and photocarcinogenesis affect DNA repair. J Invest Dermatol, 130, 1428–37.

Tang, H. (2009) Regulation and function of the melanization reaction in Drosophila. Fly, 3, 105–111.

Tsai, C. J., Loh, J. M. & Proft, T. (2016) Galleria mellonella infection models for the study of bacterial diseases and for antimicrobial drug testing. Virulence, 7, 214–29.

Uçkan, F., Er, A. & Ergin, E. (2010) Levels of encapsulation and melanization in Galleria mellonella (Lepidoptera: Pyralidae) parasitized and envenomated by Pimpla turionellae (Hymenoptera: Ichneumonidae). Journal of Applied Entomology, 134, 718–726.

Whitten, M. M. A. & Coates, C. J. (2017) Re-evaluation of insect melanogenesis research: Views from the dark side. Pigment Cell Melanoma Res, 30, 386–401.

